# Pre-training with a rational approach for antibody sequence representation

**DOI:** 10.1101/2023.01.19.524683

**Authors:** Xiangrui Gao, Changling Cao, Chenfeng He, Lipeng Lai

## Abstract

Antibodies represent a specific class of proteins produced by the adaptive immune system in response to pathogens. Mining the information embedded in antibody amino acid sequences can benefit both antibody property prediction and novel therapeutic development. Protein-specific pre-training models have been used to extract latent representations from protein sequences, structural, functional, and homologous information. However, compared to other proteins, antibodies possess unique features that should be incorporated using specifically designed training methods, so there is still room for improvement in pre-training models for antibody sequences. On the one hand, existing protein pre-training models primarily utilize language models without fully considering the differences between protein sequences and language sequences. In this study, we present a Pre-trained model of Antibody sequences trained with a Rational Approach for antibodies (PARA), which employs a strategy conforming to antibody sequence patterns and an advanced natural language processing self-encoding model structure. We demonstrate PARA’s performance on several tasks by comparing it to various published pre-training models of antibodies. The results show that PARA significantly outperforms existing models on these tasks, suggesting that PARA has an advantage in capturing antibody sequence information. We believe that the antibody latent representation provided by PARA can substantially facilitate studies in relevant areas. PARA is available at https://github.com/xtalpi-xic.

## Introduction

The adaptive immune system generates antibodies of immense diversity to specifically recognize and neutralize invading pathogens. Antibodies are Y-shaped protein complexes that contain several functional regions, including Fab, Fc, and Fv regions. Among them, V(D)J gene recombination contributes to the initial diversity in the Fv region, while antigen-specific affinity maturation drives somatic hypermutation to enhance diversity throughout the entire protein^1^. Being able to model the complexity of antibody diversity will facilitate the study of immune system as well as benefit the computer-aided antibody drug design.

Currently, self-supervised pre-training on large-scale unlabeled data enables the achievement of robust model performance. Numerous studies are presently focusing on the pre-training of protein sequences. Some research has indicated that language models trained on protein sequences can effectively capture the structural, functional, and co-evolutionary information of proteins. Recently, based on large pre-trained models of protein sequence data, researchers have developed novel protein structure prediction models that replace multiple sequence alignment in AlphaFold with protein pre-training models, yielding satisfactory results^2,3^.

There are several available pre-trained models for antibody sequence, including representative models such as AntiBERTa, AntiBERTy, and ABLang^4–6^. AntiBERTa is a model pre-trained on 57 million human antibody sequence data (42 million heavy chains and 15 million light chains), featuring a structure and hyperparameters identical to those of RoBERTa. The authors demonstrated AntiBERTa’s effectiveness by constructing a model that uses the latent representations of antibody sequences from AntiBERTa to predict the probability of a residue serving as an antigen-binding site on the antibody. AntiBERTy is another pre-trained model with a structure and hyperparameters identical to BERT, trained on 558 million antibody sequences. By obtaining antibody sequence representations from AntiBERTy, researchers can cluster antibodies into trajectories similar to affinity maturation. ABLang shares the same model structure as RoBERTa and has trained heavy chain and light chain models on the heavy and light chains of Observed Antibody Space (OAS)^7^, respectively.

In addition to showcasing the potential applications of the latent representations provided by ABLang, the authors also emphasized the model’s impressive performance in recovering missing fragments in antibodies. These works resulting in pre-trained models for antibody sequences that have achieved satisfactory performance in certain downstream tasks. However, these pre-training models were trained without fully considering the difference across antibody regions (e.g. Fv vs Fc), so we hypothesize that they can still be improved.

In this study, we propose a novel antibody pre-training model (Pre-trained model of Antibody sequences trained with a Rational Approach for antibodies, PARA, Fig. 1), which demonstrates superior performance across multiple tasks compared to existing models. Our training dataset comprises approximately 18 million human antibody sequences, including 13.5 million heavy chains and 4.5 million light chains. PARA is built upon the general DeBERTa^8^ model architecture, and was trained under a rational approach by considering the unique characteristics of antibody sequences. For example, the antibody framework (FRs) within the Fv region display limited variability, while the CDRs exhibit higher variability, shaped significantly by the antibody antigen specificity^1^. As a result, there’s a pronounced redundancy of information in the FRs of antibody sequences, whereas the CDRs, and particularly CDR3, contain more information and contribute significantly to the diversity of antibody sequences. Such regional differences of antibody sequences suggest the classic way of training natural language models might not suitable for antibody sequences. Hence, to train PARA model, we employed a combination of span masking and random word masking during the pre-training phase, with an increased probability for CDR-H3 masking (see Methods). Such training strategies emphasize more on the CDR-H3 region, which result in higher CDR-H3 and FR recovery accuracy. In the tasks for heavy-light chain pairing and antibody binding prediction, utilizing PARA’s latent representations also yields higher prediction accuracy. These observations suggest our unique rational approach for training PARA allows the model to learn more meaningful antibody representations.

**Figure 1.**
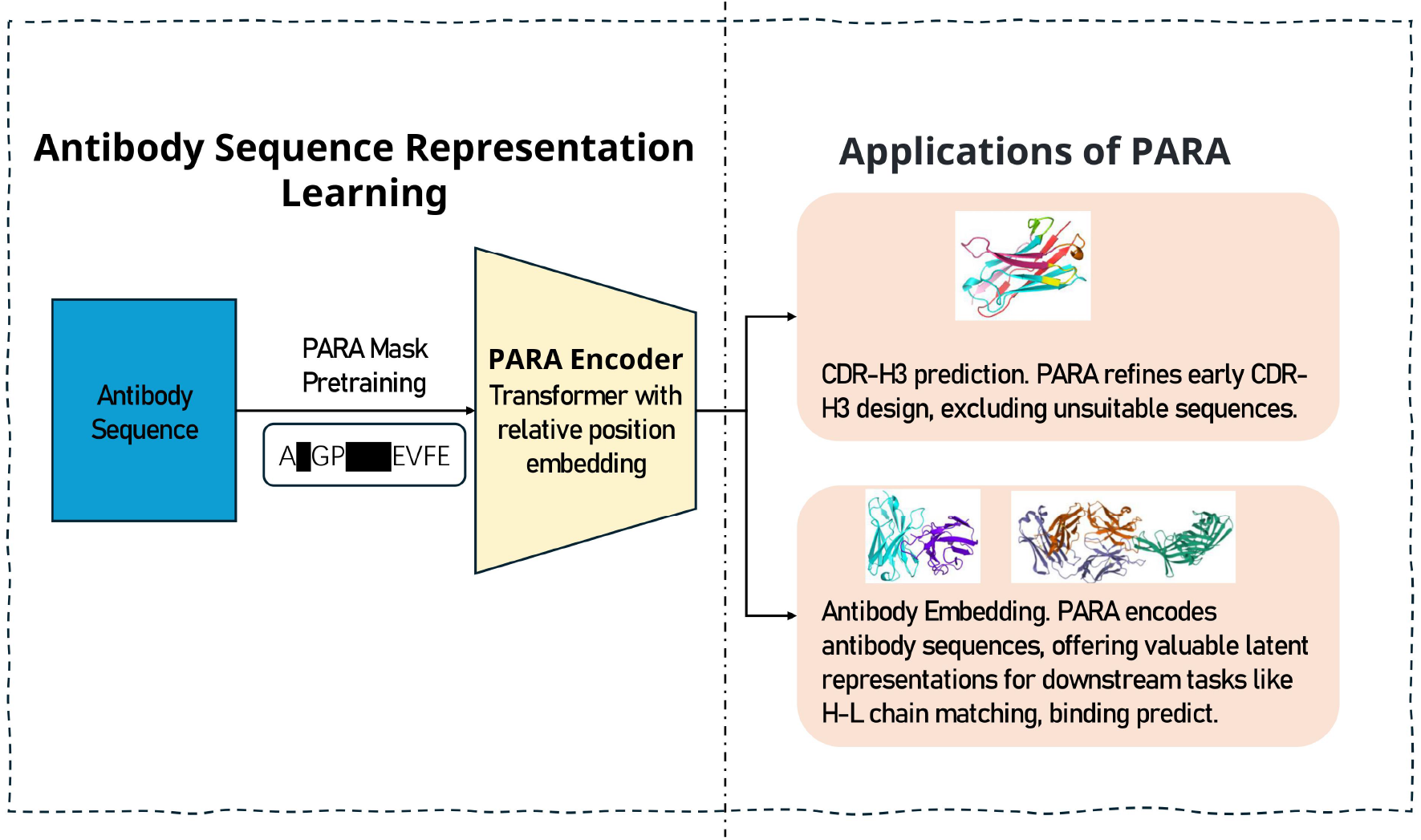
The training and prediction process of PARA. We engineered a pre-training task that is targeted at antibody sequences to yield a specialized model for antibody sequence analysis. The model serves two primary functions: firstly, it aids in predicting the masked regions of antibody sequences, which is instrumental in antibody engineering by narrowing down the vast number of potential sequence combinations. Secondly, the latent representations of antibody sequences obtained through PARA encoding can be utilized as input for downstream tasks, particularly beneficial when the dataset for such tasks is limited in size.

## Methods

### Antibody sequence processing

Our training data consist of the Fv fragments of human antibodies sequences derived from the Observed Antibody Space (OAS) dataset^7^. Within the unpaired data of the OAS dataset, we first employ Linclust to cluster the heavy and light chain sequences with 80% sequence identity^9^. This clustering step is crucial as it aims to eliminate highly similar antibody sequences, thereby enhancing the quality of our training data and reducing the volume of training material in a meaningful way. Numerous studies have highlighted that the removal of low-quality and highly redundant data can significantly improve the training process^6,10,11^. Subsequently, we retain clusters with multiple members and select the longest sequence as the representative for each cluster. To fully utilize the data, we re-cluster single-member clusters at 50% sequence identity and retain clusters with multiple members once again. The selected sequences are then randomly divided into training, validation, and test sets. The training set comprises 13 million heavy chain sequences and 4 million light chain sequences, the validation and test set both include 175,000 heavy chain sequences and 57,000 light chain sequences.

To further enhance the robustness of our model, we have adopted a data augmentation that involves the random truncation of 0 to 3 amino acids from the N-terminus and C-terminus of the antibody sequences. This trick is analogous to the common practice in natural language processing where language sequences are randomly truncated to improve the model’s ability to handle variable-length inputs and to prevent overfitting to sequence.

For tokenizing and encoding the sequences, we added two special tokens, [CLS] and [SEP], at the beginning and end of the sequence, respectively. For the sake of convenience, we did not train the heavy chain sequences and light chain sequences separately like ABLang, nor did we add corresponding special tokens to deliberately mark their differences.

### The PARA Architecture

Our PARA model is grounded in the DeBERTa^8^ architecture, a variant within the transformer model family, which forms the structural foundation for our antibody sequence modeling task. The transformer framework is adept at processing sequential data, primarily through its self-attention and feed-forward network components. DeBERTa enhances this framework with a disentangled attention mechanism that effectively integrates the relative positional relationships and categorical information of each token in a sequence, thereby augmenting the model’s capabilities.

#### Transformer Architecture

In our work, we construct a neural network architecture using multiple layers of Transformer units to form an encoder. This encoder processes the input sequences, leveraging the self-attention mechanisms within the Transformer layers to capture the intricate dependencies among the data. The encoded representations are then fed into a linear layer, which serves as the final step in our model to predict the categories of the masked tokens.

The attention mechanism in the Transformer model is defined as follows:

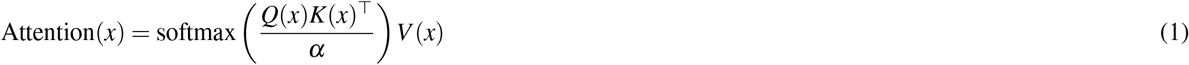

where *Q*(*x*), *K*(*x*), and *V* (*x*) are distinct representations of each token’s latent information within the sequence, obtained through separate linear transformations. These transformations are initially derived from the token embeddings. Notably, additional details such as relative positional encodings can be integrated into *Q, K*, and *V* to enhance the model’s performance. The scaling factor *α*, represented by *α*, is used to stabilize the gradients during training.

The feed-forward network (FFN) is an essential component of the Transformer model. It consists of two linear transformations separated by a non-linear activation function. The FFN can be defined as:

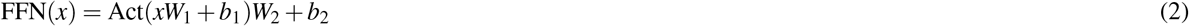

where *x* represents the input to the FFN, Act denotes activation function, and *W*_1_, *b*_1_, *W*_2_, and *b*_2_ are learnable parameters (weight matrices and bias vectors).

To sum up, the Transformer layer architecture includes the attention mechanism and the feed-forward network together with the input embedding, which can express as the following operations:

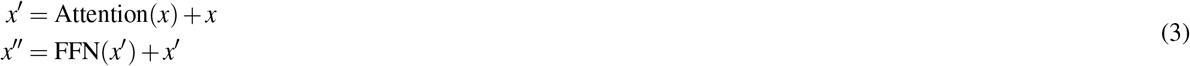

where *x*^*′*^ is the transformation after considering the attention mechanism, which serve as input to the FFN and output *x*^*′′*^.

#### Disentangled Self-Attention with Relative Position Encoding (DeBERTa)

In sequence-based neural network models, position encoding is essential for providing the model with information about the order of elements. There are two main types of position encoding: absolute and relative.

Absolute Position Encoding assigns a unique identifier to each position in a sequence, allowing the model to recognize the order of inputs. This method is simple and effective but can be limited by fixed sequence lengths and may not generalize well to longer sequences not seen during training^12^.

Relative Position Encoding encodes the position of tokens relative to each other, focusing on the distance between pairs of tokens. This method is more flexible, as it is not bound to fixed positions and can handle variable sequence lengths. It is particularly useful for tasks where the relationship between tokens is more important than their individual positions^13^.

In DeBERTa, the self-attention mechanism is augmented to encode more precise positional relationships between tokens. This enhancement is realized by disentangling the attention computation into distinct streams: ‘content-to-content’, ‘content-to-position’, and ‘position-to-content’ interactions. Specifically, the model encodes each token using two vectors—a ‘content vector’, which captures the semantic meaning of the token (essentially the token’s representation in the sequence), and a ‘position vector’, which encodes the token’s position within the sequence. The attention mechanism then calculates scores that integrate this semantic content with their relative positions, allowing the model to maintain a nuanced understanding of both the meaning of each token and its context within the sequence.

Specifically, the disentangled self-attention mechanism for a single head can be expressed as follows:

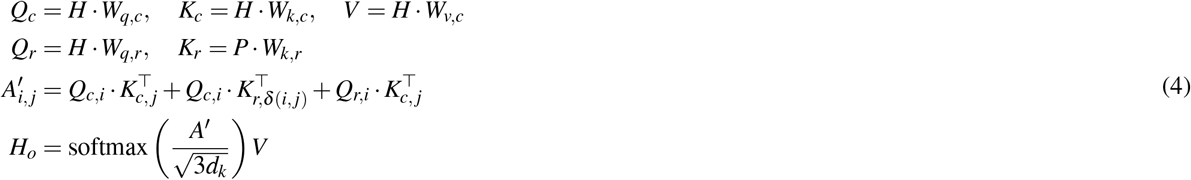

Here, *H* ∈ ℝ^*N×d*^ denotes the input hidden vectors, *H*_*o*_ ∈ ℝ^*N×d*^ is the output of self-attention, and *W*_*q,c*_,*W*_*k,c*_,*W*_*v,c*_,*W*_*q,r*_, 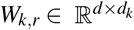 are the projection matrices for the content’s query, key, and value, and for the relative positions’ query and key, respectively. *A*^*′*^ ∈ ℝ^*N×N*^ is the attention score matrix, *N* is the sequence length, *d* is the dimension of the hidden states, and *d*_*k*_ is the dimension of the key vectors. The relative position embeddings are represented by the matrix *P* ∈ ℝ^(2*k*+1)*×d*^, where each row corresponds to a relative position embedding vector. The relative position index *δ* (*i, j*) is computed as:

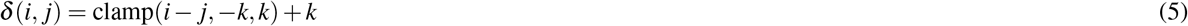

where clamp(*x, min, max*) is a function that limits the value of *x* to the range [*min, max*]. The term *k* is a hyperparameter(in this work k=128) that defines the maximum relative distance considered by the model. This clamping ensures that the relative positions are represented within a fixed range, allowing the model to generalize to sequences with length beyond those in the training set.

The attention scores *A*^*′*^ are scaled by 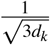 before applying the softmax function to normalize them. This scaling factor is used to stabilize gradients during training, as it accounts for the increased variance introduced by the summation of three different dot products in the attention computation.

DeBERTa effectively models the interplay between content and position by focusing on these key interactions, leading to its strong performance on language understanding tasks (For further details, refer to the discussion section). Absolute position encoding and relative position encoding utilize the same underlying information, yet relative position encoding more closely resembles the antibody sequence numbering system we employ.

### Pretraining Strategy for Enhanced PARA Performance

#### High proportion of masking

Given the limited variation in the conserved region of antibody sequences, the model can readily predict the masked portions. However, when difficulty of the pre-training task is too low, the training level becomes insufficient, hindering the model from achieving optimal performance. Conversely, if the pre-trained data contains less but valid information, the model will strive to explore patterns, resulting in a more robust feature extraction and generalization capability^14–16^. In natural language tasks, mask language model(MLM)-based self-encoding models like BERT and T5 typically employ a 15% masking rate, a setting followed by most self-encoding models^17^.

Recent studies, however, have demonstrated that a larger masking rate, combined with a larger model and learning rate, may yield better training outcomes of the final model^18^. In computer vision (CV), Mask Image Models (MIM) such as MAE, models adopt substantially higher masking rates (70%-90%), outperform those with lower masking rates^19^.

The crux lies in the amount of effective information present in the data. Compared to text, which is highly condensed and semantically rich, images are less semantically dense and contain a larger volume of information that may not be directly relevant to the task at hand, such as classification. This excess of information in images, much of which does not contribute to the understanding required for the classification task, constitutes redundant information. This redundancy can impede th model’s ability to efficiently learn key features, as it has to sift through and process a vast amount of extraneous data to identify the salient features that are truly informative for the task. In antibody sequences, numerous residues are conserved, exhibiting minimal variation and a strong correlation with their absolute positions in the sequence. If only 15% of the region is masked, domain experts with a solid understanding of antibody sequences can effectively predict the masked region, especially within the conserved areas, through sequence alignment. For models with a large number of parameters, performing MLM tasks using this information is relatively straightforward. Nonetheless, it is desirable for the model to uncover the latent information between residues in the sequence to predict the types of residues at the masked positions.

Based on the aforementioned analysis, we trained the PARA model using several different masking rates, 15%, 30%, 50%, 70% and 90%, to evaluate their respective performances.

#### PARA Mask

The PEGASUS framework employs a unique pre-training strategy to enhance model performance in text summarization tasks^20^. Unlike the random token masking approach in MLM tasks, PEGASUS masks the first, last, and central sentences with the highest correlation to other sentences, significantly improving model performance in text summarization tasks. In the antibody Fv region, CDRs exhibit high variability, particularly in CDR-H3. Our goal is to use targeted masking of CDR-H3 to enhance the model’s ability to capture high-quality CDR-H3 features.

In many NLP tasks, models trained with Span Mask outperform those trained with random masking. Span Mask ensures language continuity, reasonably increases language model learning difficulty, and encourages deeper exploration of text relationships. The antibody Fv region sequence comprises distinct partitions, including FRs and CDRs. We believe the span mask method is better suited for antibody pre-training.

Based on this, we designed a novel antibody masking strategy called PARA mask. For the heavy chain, we initially set a 70% mask rate, masking part of CDR-H3. We then use span mask to randomly mask remaining regions until the predetermined total masking rate is achieved. For the light chain, we apply a mask rate and span mask without targeting specific regions, since the light chain sequences are generally more conserved. Fig. 2 shows masked areas in cyan, comparing PARA mask with random mask strategy and span mask. We propose that variations in the conserved regions of the antibody sequence, such as deletions and misalignments, should not impact the embedding of CDRs.

**Figure 2.**
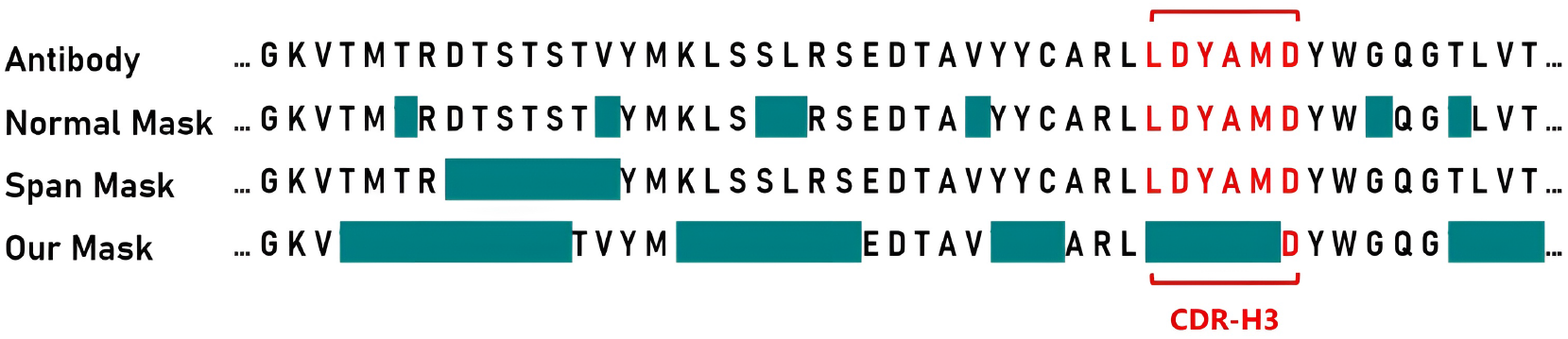
Masking strategies for Antibody sequences. For presentation purposes, we have isolated the segment of the antibody heavy chain Fv region that includes the CDR-H3, indicated in red. The cyan color signifies the areas that will be masked during pre-training. With the PARA approach, we first mask portions of the CDR-H3, and then apply span masking to other regions.

#### Pre-training setting

We trained the model using the PARA mask language model method. Cosine learning rate decay and warm-up were employed, and label smoothing technique was applied when calculating cross-entropy, with the corresponding hyperparameter set to 0.1. Our model was trained on a V100 with 4 cards and 32GB of memory for two weeks. For detailed training configurations, refer to Table 1.

**Table 1.** PARA training configurations.

| Config | Value |
| --- | --- |
| Optimizer | AdamW |
| Base Learning Rate | 3e-4 |
| optimizer momentum | $\beta_1, \beta_2 = 0.9, 0.99$ |
| Weight Decay | 0.01 |
| Batch Size | 784 |
| Learning Rate Schedule | Cosine decay with warmup |
| Warmup epochs | 15 |
| Training Epochs | 200 |
| Label Smoothing | 0.1 |
| Mask Ratio | 15%, 30%, 50%, 70%, 90% |
| Augmentation | Random truncation |

### Methods for Heavy and Light Chain Matching with PARA

To assess the capability of the PARA model in providing potential representations of antibody sequences, we applied it as an encoder for various downstream tasks. In this study, we extracted the latent representations of the heavy and light chains using PARA and performed heavy chain light chain pairing tasks based on these representations.

Accurate light and heavy chain pairing information is crucial for antibody research, as different H-L chain combinations affect the folding, expression, and antigen-binding characteristics of antibodies^21^. We adopted a contrastive learning approach, using existing antibody pairing data in OAS to train our H-L chain pairing model that can discern the likelihood of a given H-L chain pairing occurring in a natural environment.

Specifically, in the data preparation phase, we clustered approximately 200,000 heavy-light chain pair data in the OAS database with a threshold of 0.9 and selected the cluster centers, ultimately obtaining about 40,000 unique antibody sequences. Based on the clustering results, we randomly selected 90% of the clusters and their members as the training dataset, with the remainder serving as the test dataset.

For the training process, positive samples consist of matched H-L chains, while negative samples are generated by randomly pairing the heavy chain from the positive samples with light chains from OAS that have less than 85% similarity. We used a pre-trained antibody sequence model to extract the latent representations of the heavy and light chains, then averaged these representations along the sequence dimension to obtain representations for each chain. These representations were processed through a dedicated projection layer containing a two-layer multilayer perceptron (MLP) to map the latent representations of the heavy and light chains to the same feature space (Fig. 3a). Subsequently, we normalized both latent vectors using L2 normalization and calculated the cosine similarity between them, standardizing this similarity to the [0, 1] range. During the training process, we froze the parameters of the pre-trained PARA model and only trained the projection layer to focus on learning the similarity between representations.

**Figure 3.**
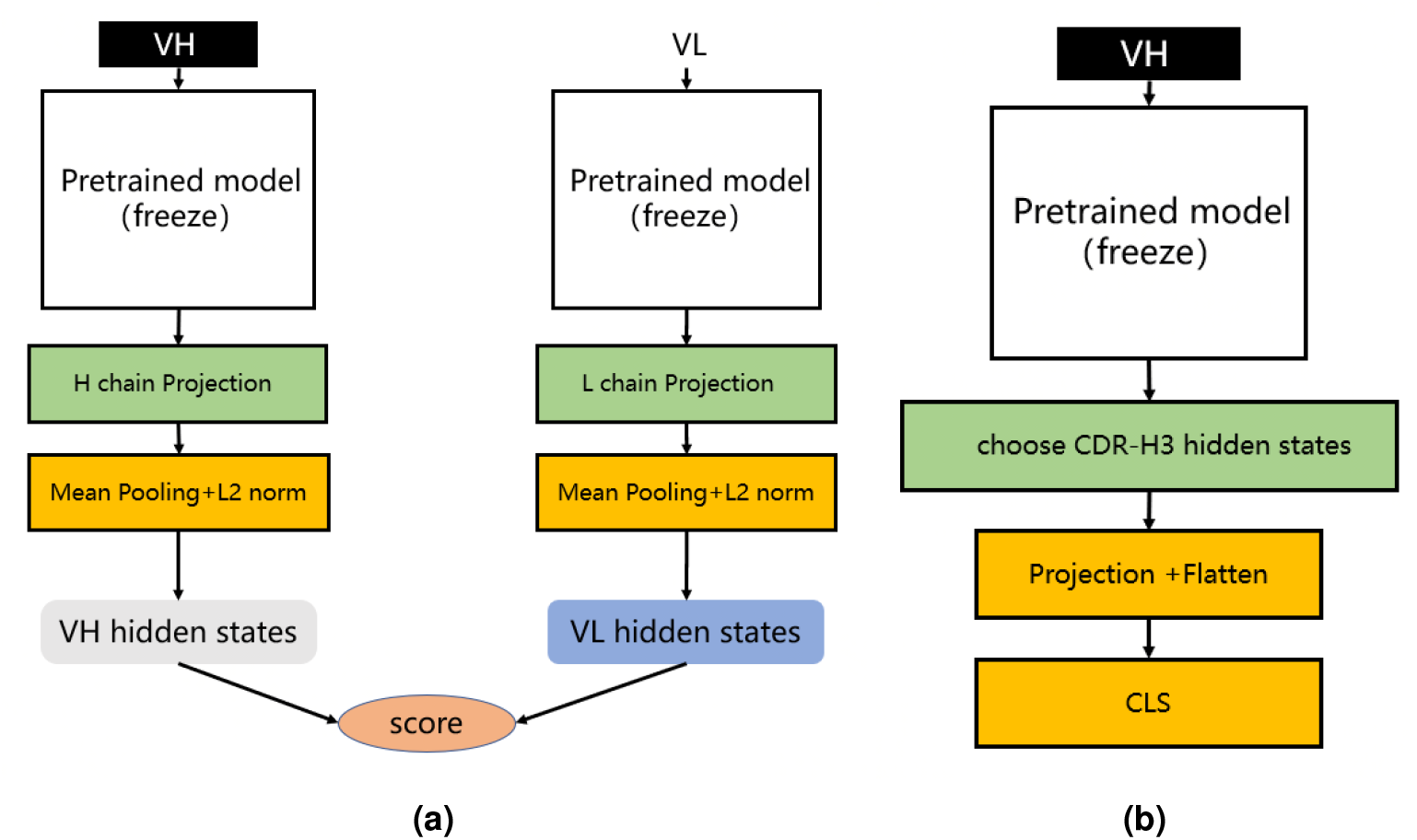
(a) Heavy and light chain matching model. We utilize different pre-trained models to extract the latent representations of heavy and light chains. During training, each training sample consists of one positive sample and multiple negative samples. We calculate their matching scores separately and compute the loss using infoNCE loss. (b) Antibody binding prediction model. We designed an antibody affinity prediction model, as illustrated in the figure. Firstly, we employed a pre-trained deep learning model to extract the latent representation of the antibody VH sequence. Subsequently, we selected only the latent representation of the CDR-H3 and reduced its dimensionality and flattened it using a single-layer multilayer perceptron (MLP) projection layer. Finally, this representation was fed into a two-layer MLP classification layer for a binary classification task, where antibodies with affinity were labeled as 1 and those without affinity were labeled as 0. We used cross-entropy as the loss function to optimize the model.

### Methods for Antibody Binding Prediction with PARA

We divided the HER2 dataset into 15,098 training samples, 3,236 validation samples, and 3,235 test samples, using the entire VH sequence as our input. To ensure the reliability of our findings, we divided the dataset into five separate parts using different random seeds and calculated the average of the performance metrics across these five distinct splits for each model to determine the final results.

## Results

First, we compared PARA with AntiBERTy and ABLang to evaluate their prediction capabilities in the CDR-H3 and FRs. We then used PARA as a sequence encoder to extract antibody sequence representions for antibody light chain and heavy chain pairing tasks and antibody binding prediction tasks.

### Ablation Studies on PARA Model for CDR-H3 Restoration

In our study, we leveraged a pre-trained model designed for masked data tasks. To avoid data leakage in our experiments, we curated a test set by randomly selecting 90,096 antibody heavy chain sequences from the model testing set, ensuring these sequences were not included in the training dataset used to train PARA. We also considered augmenting our test set with newly available antibody sequences from public databases, such as the Structural Antibody Database (SAbDab), focusing on data published in 2022 or later. This approach aimed to minimize the likelihood of the task model having previously encountered the data. However, the scarcity of such data, compounded by further reductions post-similarity-based clustering, would not permit a comprehensive evaluation of the model’s performance. Consequently, we opted to utilize the pre-established test set, confirming that PARA had no prior exposure to these sequences.

The primary goal of our research was to pre-train a model adept at extracting robust latent representations of antibody sequences, with an emphasis on the CDR-H3 region. To achieve this, we conducted ablation studies on the PARA model’s design choices within the CDR-H3 restoration task, focusing on two main aspects: the effect of different masking strategies and the role of positional embeddings.

In our masking strategy ablation studies, we considered two key factors: the sequence masking rate and the targeted masking of the CDR-H3 region. We experimented masking rates of 15%, 30%, 50%, 70%, and 90%. The model’s performance was evaluated using top-1 and top-3 prediction accuracies. Notably, for the 15% masking rate, we employed the settings used in BERT, which are commonly adopted in NLP for masked language modeling tasks.

The results of these experiments are presented in Fig. 4a, an asterisk (*) following a masking rate denotes the use of a uniform random masking method rather than the PARA mask method. We observed that increasing the masking rate from 15% to 70% improved the model’s ability to restore CDR-H3, aligning with our initial hypothesis that a reasonable increase in task difficulty enhances predictive capabilities. However, at a 90% masking rate, the model’s performance declined, likely due to an excessive portion of the sequence being masked, resulting in insufficient information. Comparing the mask-50 and mask-50* experiments, we found that targeted masking of CDR-H3 significantly improved restoration rates. Yet, as the masking rate increased further, this improvement diminished, as seen in the comparison between mask-70 and mask-70*. We believe this is due to a high masking ratio increasing the likelihood of CDR-H3 being masked. Based on these findings, we selected the mask-70 setting as the default for subsequent use with PARA.

**Figure 4.**
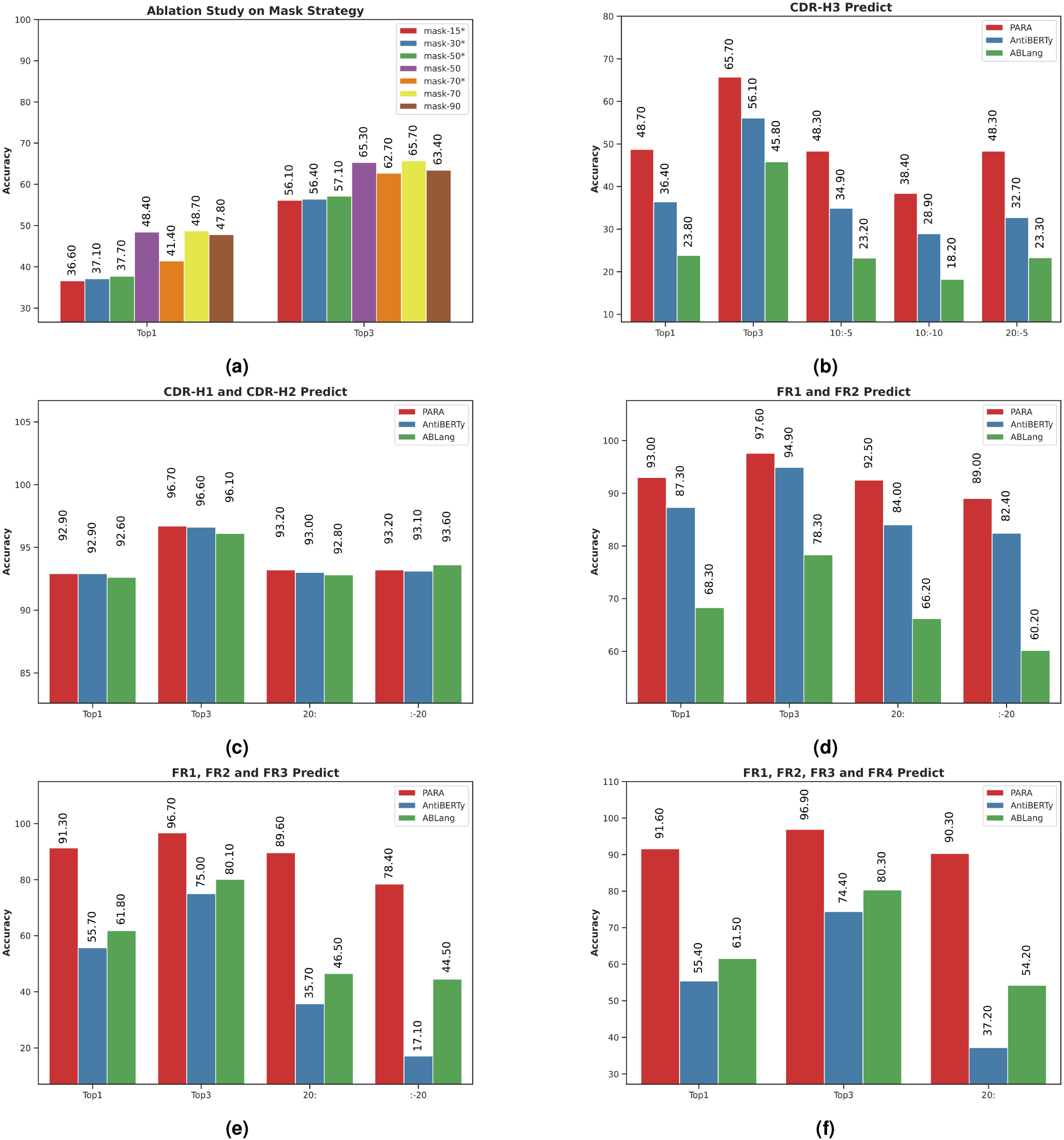
Model performance in antibody sequence prediction. Top1 and Top3 denote the highest accuracy and the average of the top three accuracies, respectively. Specific subsequences of the antibody sequences were masked to test model robustness, with Top1 accuracy compared using slice notation, e.g., “10:-5” for masking from the 10th to the 5th position from the end. (a) PARA Mask ablation study evaluated recovery accuracy in the CDR3 region, with “*” indicating random masking and no “*” indicating PARA Mask. PARA 70 was selected as the standard configuration after showing superior performance at various masking ratios. (b) PARA outperformed other open-source models AntiBERTy and ABLang in CDR-H3 masking prediction, maintaining high accuracy even with truncated sequences. (c) All models showed high Top-1 and Top-3 accuracies in CDR-H1 and CDR-H2 regions, despite sequence truncation. (d) PARA and AntiBERTy outperformed ABLang in FR1 and FR2 region masking prediction. (e) PARA excelled in masking prediction accuracy in FR1, FR2, and FR3 regions. (f) PARA maintained high accuracy in FR1, FR2, FR3, and FR4 regions, even with extensive sequence masking, outperforming other models. PARA consistently achieved the best results in experiments (b) to (f), demonstrating optimal and stable performance under sequence truncation or extensive masking, unlike the significant performance decline in other models.

For the positional embedding ablation studies, we utilized a 70% masking rate with targeted CDR-H3 masking. We compared the use of absolute positional encoding (Abs.Pos.Enc.) and the relative positional encoding (Rel.Pos.Enc.) from DeBERTa. To further assess the impact of different positional encodings, we truncated the antibody sequences at various points, with notations such as 10:-5, 10:-10, and 20:-5 indicating the start and end points for truncation before predicting the masked CDR-H3. The experiments (Table 2) demonstrated that relative positional encoding improved the model’s performance even with only CDR-H3 masked. When sequences were truncated, models with absolute positional encoding experienced a significant drop in predictive capability, whereas those with relative positional encoding were minimally affected. This suggests that relative positional encoding allows the model to better learn the interactions between amino acids rather than rigidly memorizing the distribution at absolute positions. In machine learning, overly strong features can prevent a model from being adequately trained.

**Table 2.** Results of the ablation study on positional embeddings with PARA70.

| CDR-H3 | Top1 | Top3 | 10:-5 | 10:-10 | 20:-5 |
| --- | --- | --- | --- | --- | --- |
| Abs.Pos.Emb. | 46.8 | 64.3 | 38.2 | 31.4 | 41.7 |
| Rel.Pos.Emb. | 48.7 | 65.7 | 48.3 | 38.4 | 48.3 |

### PARA demonstrates a strong robustness with a high CDR restoration capacity

We compared the ability of different models to recover CDR-H3. Fig. 4b shows the prediction accuracy of the three models, PARA, ABLang, and AntiBERTy. PARA represents the model mask-70 in Fig. 4a. Top1 and Top3 refer to the Top1 and Top3 accuracies.

It can be observed that when CDR-H3 is completely masked, PARA has the highest accuracy in restoring it. When the antibody sequence is truncated, the performance of all models decreases, but the change in PARA is the smallest, with a significant decrease only at 10:-10, and a decrease of only 0.4% at 10:-5 and 20:-5. The performance of the ABLang and AntiBERTy models decreases significantly when the sequence is truncated.

Furthermore, we tested the masking of CDR-H1, CDR-H2, and the simultaneous masking of both CDR-H1 and CDR-H2 (Fig. 4c). The masking rate was approximately 8% for individual CDR-H1 or CDR-H2, and 16% when both were masked simultaneously. The results were similar across these tests, so we only present the findings for the simultaneous masking of CDR-H1 and CDR-H2. Notably, the predictions for CDR-H1 and CDR-H2 across the three models were very close, likely due to the relatively small variations in these regions.

Although the pre-trained models have good memory and capability for generalization, the predicted results still differ from the true distribution. Here, we compare the distribution of amino acids in the CDR-H3 with the amino acids distribution predictions made by other pre-trained models, and find that the distribution given by PARA is closer to the true distribution (Fig. 5).

**Figure 5.**
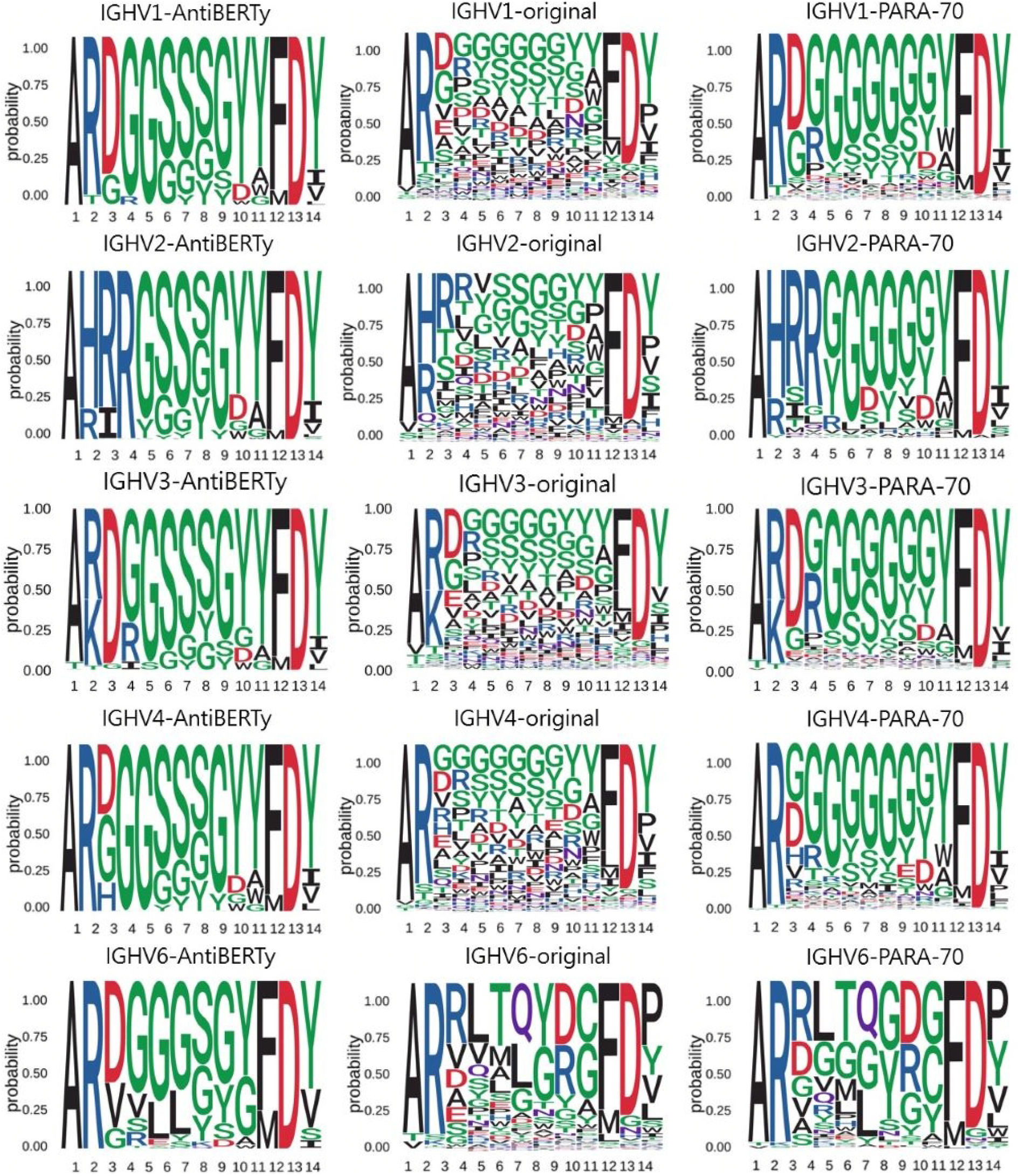
Amino acid distribution in CDR-H3. We obtained distributions from the CDR-H3 of the real antibody sequences, the CDR-H3 of the AntiBERTy predicted sequences, and the CDR-H3 of the PARA predicted sequences. In this test, we focused on the diversity of predictions. We tested IGHV1, IGHV2, IGHV3, IGHV4, IGHV5, IGHV6 and IGHV7 and present some of the results. Using data from the test set, we found that the CDR-H3 distributions derived from PARA are closer to the original distribution.

We believe that by increasing the masking rate, PARA training can infer most of the information of the entire sequence with only a small amount of input information. From another perspective, the masked sequence can be seen as a compression of the information in the complete sequence, making it less susceptible to the issue of information redundancy. Excessive redundancy may lead to overfitting during model training, resulting in more uniform output results.

We then compared the amino acid distribution in the CDR-H3 region as predicted by PARA and AntiBERTy with the actual distribution. Here, we focused on these two methods because they demonstrated comparable performance, both significantly surpassing ABLang, in antibody sequence prediction (Fig. 4b). In order to eliminate the impact of different models’ prediction accuracy on the prediction distribution, we deliberately sampled from the test set to ensure that the accuracy of the two models in the sampled samples is comparable. To brief the process, we divide the heavy chains in the test set by the IGHV gene and CDR-H3 length. We statistically obtained the true distribution of amino acids in the CDR-H3 for the same V gene and the same CDR-H3 length. We obtained two other distributions from the prediction results of the statistical model.We obtained two other distributions from the prediction results of the statistical model and show that the predicted result of PARA, unlike AntiBERTy, is closer to the real antibody sequence (Fig. 5).

### PARA demonstrates a strong robustness with a high FR restoration capacity

In developing PARA’s training strategy, we implemented span masking on the CDR-H3 region to bolster the model’s proficiency in processing highly variable regions. This approach, however, prompted a concern regarding its potential impact on the model’s accuracy in predicting the more conserved FRs. Given the significant conservation of FRs, experts can typically deduce the masked segments within these regions through sequence alignment with notable accuracy. Our goal was for PARA to demonstrate a similar, if not superior, level of accuracy.

To evaluate the model’s resilience, we conducted tests where we selectively masked FR1 and FR2 and truncated the sequences. PARA consistently maintained accuracy across these modifications, outshining the counterparts. The minimal impact of targeted masking and sequence truncation on PARA’s performance is substantiated by the data in Fig. 4d, 4e, and 4f.

For a more streamlined analysis, we limited our testing to two scenarios: truncating either the first 20 or the last 20 residues of the sequence. We excluded the truncation of the last 20 residues for FR4, given its position at the end of the FV sequence. These tests lend further support to the notion that antibody sequences exhibit significant redundant information, and that a modest masking rate might be insufficient for thorough model training. Moreover, an expert with knowledge of antibody sequences could likely reconstruct the conserved regions in FV that have been masked, using sequence alignment.

It is particularly noteworthy that PARA’s performance remained robust across varying sizes of masked FRs, whereas the performance of AntiBERTy and ABLang deteriorated as the masked region expanded. Despite an increased masking of FR4, there was no significant decline in the model’s performance, underscoring PARA’s exceptional ability to handle sequence alterations within conserved regions (Fig. 4e and Fig. 4f).

### Results of Heavy and Light Chain Matching Model with PARA

We then posed a question: Given a heavy chain H1 and two light chains (L1, L2) with sequence information and H1 is known to pair with L1, can the model accurately identify which light chain more commonly pairs with H1 when the sequence similarity between L1 and L2 is low?

To answer this question, we conducted a heavy-light chain pairing task using three different pre-trained language models as backbones and tested the Area Under the Receiver Operating Characteristic (auROC) and Area Under the Precision-Recall Curve (auPRC) on the test set. As shown in Fig. 6, the experiment results indicate that the heavy-light chain pairing model trained with PARA outperforms those trained with AntiBERTy and ABLang in terms of auROC and auPRC on the test set (with the AUC for random matching being 0.5). Specifically, Fig. 6 illustrates that PARA achieves higher auROC and auPRC scores compared to AntiBERTy and ABLang, indicating better performance in predicting correct heavy-light chain pairs. The “Chance” line in the auROC plot indicates the baseline performance of a random classifier, with an AUC of 0.5, signifying that any predictive accuracy above this line is better than random guessing. Since the model and training parameters other than the antibody latent representations are consistent, we believe that PARA’s latent representations may contain more information about the antibody sequences. Interestingly, ABLang performs poorly in this task. Unlike AntiBERTy and PARA, ABLang models the heavy and light chains independently, which may fail to capture the pairing information coded within the antibody heavy and light chain sequences.

**Figure 6.**
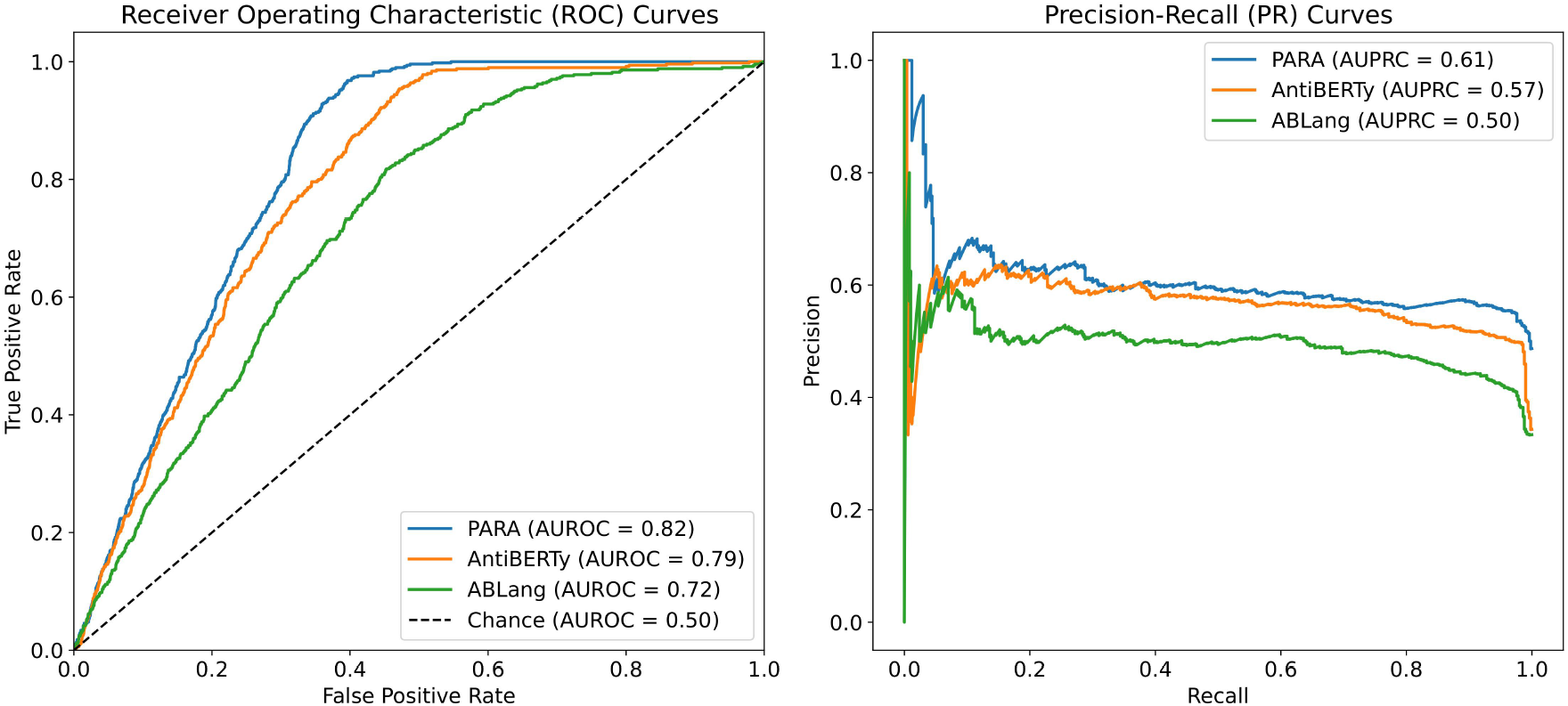
auROC and auPRC for Heavy Light Chains Matching. We trained AntiBERTy and PARA as antibody sequences encoder on OAS antibody paired sequences with contrast learning, respectively. The matching model obtained by PARA performs better with the same training set and training method. The “Chance” line in the auROC plot indicates the baseline performance of a random classifier, with an AUC of 0.5, signifying that any predictive accuracy above this line is better than random guessing.

To mitigate potential biases caused by randomness, we performed five repeated validations, each time randomly selecting negative samples and then calculating the average auROC value. We also included randomly matched heavy and light chains as a control group for comparison.

## Results of Antibody Binding Prediction with PARA

Antibodies function by binding antigens, and being able to predict their binding specificity would greatly benefit the downstream application of the trained large language models. To evaluate this, we utilized the three pre-trained models to train a binding prediction model. Mason et al.^22^ employed a deep learning approach to predict antigen specificity from different antibody sequences. Here, we use their HER2 antibody data to compare PARA, AntiBERTy, and ABLang as antibody sequence encoders. These encoders were connected to a projection layer and classification layer composed of an MLP to predict whether the antibody binds to HER2 or not (Fig. 3b). The results of the experiments are presented in Fig 7a. Specifically, we used the best-performing model on the validation set to make predictions on the test set and calculated the scores for the test set. The PARA model outperformed other models on both the test and validation datasets.

**Figure 7.**
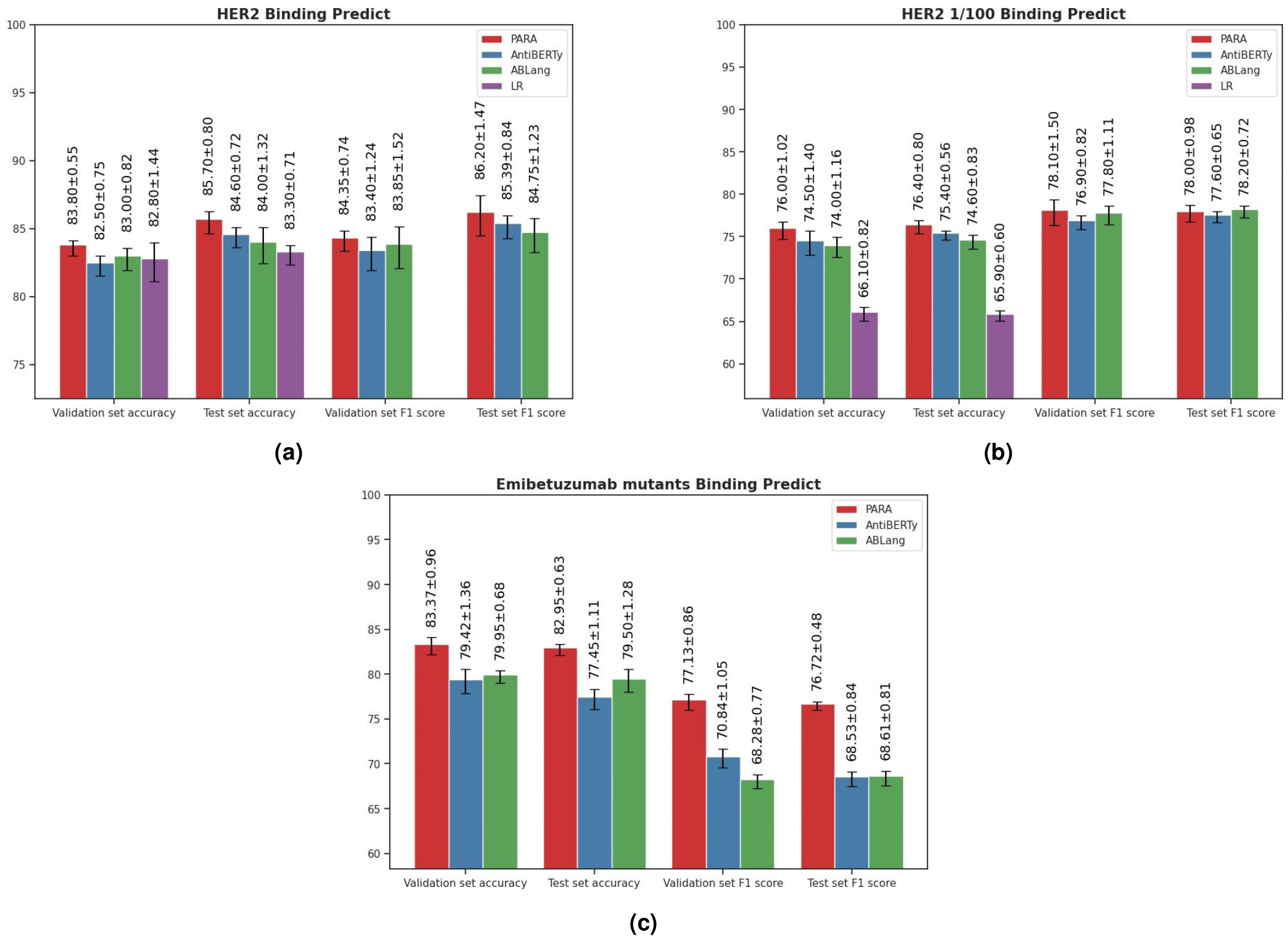
**a)** Results of Models on HER2 Antibody Dataset. The logistic regression (LR) benchmark is derived from the methodology of Mason et al. To ensure the robustness of our model evaluation, we employed a repeated random sub-sampling validation approach, where the data was randomly partitioned into training, testing, and validation sets five separate times. The results tabulated represent the mean performance metrics across these five iterations. In addition to accuracy, we computed recall and precision metrics to provide a comprehensive analysis of the predictive capabilities of each pre-trained model under investigation. **b)** Results of Models on HER2 Antibody Dataset. HER2 1/100 signifies training with only the top 1% of the original dataset to compare pre-trained models with a baseline model initiated from scratch in a data-constrained setting. The figure presents the average validation and test accuracy (Acc), recall, and precision metrics, calculated over five repeated validation trials, for each model. **c)** Results of Models on Emibetuzumab mutants Dataset. The figure presents the average validation and test accuracy (Acc), recall, and precision metrics, calculated over five repeated validation trials, for each model.

Importantly, although the binding prediction models obtained using pre-trained models perform slightly better than the LR model used in the original study, the difference is not significant. However, pre-trained models generally perform better when data size is limited. To evaluate the robustness of these models with limited training data, we sub-sampled the training set size by randomly selecting 150 antibodies (sampling rate: 0.01) as the training set. We then conducted the same experiments and the results (Fig 7b) indicate that the pre-trained models outperform the original model, with PARA showing the strongest generalization capability on this reduced dataset.

Additionally, we included another dataset from Makowski et al.^23^, which involved generating mutants of Emibetuzumab, resulting in over 4000 antibody sequences. For this dataset, we randomly selected 100 antibody sequences as the training set, 1900 sequences as the validation set, and 2000 sequences as the test set. The purpose of selecting 100 sequences as the training set is to evaluate the generalization capability of the models. Similar to the HER2 antibody data, we used the best-performing model on the validation set to make predictions on the test set and calculated the scores for the test set. The results (Fig 7c) show that the PARA model outperformed other models on both the test and validation datasets.

## Discussion

Antibody sequences manifest unique properties when compared with natural language sequences. Languages comprise an extensive lexicon, whereas antibody sequences are constructed from a canonical set of 20 amino acids. Linguistic sequences are versatile, with various permutations conveying identical meanings. In contrast, antibody sequences are comparatively uniform, each consisting of conserved and variable regions. We hypothesize that antibody sequences are akin to pixelated image patches, as both encompass redundant information and demonstrate pronounced regional attributes. Recent research, including those on Vision Transformers, has indicated that high-ratio mask pretraining on images augments model proficiency^19,24^. Our comparative analysis of different models’ capacity to reconstruct antibodies’ CDRs and FRs suggests that our rational masking strategy is more apt for antibody sequences than conventional approaches.

In antibody sequence analysis, the CDRs, especially CDR-H3, is responsible for recognizing specific antigens, which necessitates a highly variable sequence. The sequence variability of CDR-H3 is much greater than that of the FRs. To effectively train models to predict such variable regions, we applied a high-ratio masking strategy to CDR-H3 during the learning process. This method not only increases the prediction challenge for this region but also minimizes the model’s susceptibility to potential biases in the training data. Learning from difficult samples is crucial for model convergence, a fact supported by extensive literature^14–16^. Our targeted masking provides the model with numerous challenging samples that are both reasonable and consistent with antibody sequence patterns. As a result of this targeted training strategy, our model delivers accurate predictions for both the CDRs and FRs.

The selection of position encoding in transformer models tailored for antibody sequences is crucial. Both absolute and relative position encodings convey residue order and spacing, yet they instill divergent biases affecting model behavior. Absolute position encoding, within a consistent numbering framework, aligns with the static locations in an antibody sequence. However, it may inadvertently predispose the model to rely on fixed position residue distributions, potentially curtailing its ability to generalize to sequences with non-standard numbering. Conversely, relative position encoding emphasizes the inter-residue relationships, independent of their absolute positions. This method is congruent with the intrinsic variability of antibody sequences, where the functional significance of residues is predominantly determined by their relative, not absolute, positions. It empowers the model to deduce the types of masked amino acids more efficaciously by capitalizing on the contextual interactions between residues. Furthermore, relative position encoding inherently possesses advantages such as not being constrained by sequence length, which naturally aligns with the variable lengths encountered in antibody sequences. This enhances the model’s resilience and generalization prowess. Human experts are capable of predicting the types of residues in antibody sequences even when segments are missing from the N-terminus, C-terminus, or even the middle regions. This is because experts can infer which positions are missing by analyzing the relationships between amino acids and aligning the sequence to the correct numbering. From this perspective, relative position encoding is more akin to human behavior and more rational than absolute position encoding.

It is imperative to acknowledge the rationality and value of established antibody numbering systems for sequence analysis. The concern is not with the numbering system per se, but with the model’s dependency on it during training. Our objective is to cultivate a model that is robust, adaptable, and adept at encapsulating the quintessence of antibody sequences, which is more effectively realized through relative position encoding. This encoding strategy ensures the model’s efficacy when confronted with sequences that diverge from the anticipated framework, thereby offering a more encompassing grasp of antibody sequence variability.

The *in silico* prediction of antibody binding specificity has great potential for drug design as well as vaccine development, which can lead to significant time and cost savings while greatly reducing the risk associated with downstream clinical applications. In the future, PARA will be trained with larger models and more data. We will also further optimize the training strategies by considering both antibody sequence and structure characteristics, providing latent representations that better align with the data patterns in biology We believe PARA’s ability to capture more meaningful antibody representations would benefit the future improvement of computer-aided drug design.

## Conclusion

PARA is a model pre-training method specifically designed for antibody sequences, utilizing a Transformer-based architecture and employing a high mask ratio and span mask for MLM. We conducted ablation experiments on the adopted strategies and compared PARA with other existing open-source antibody pre-training models across several tasks. The results demonstrate that the latent representations generated by PARA can better capture the information embedded within antibody sequences and exhibit robustness. We attribute this to PARA’s training strategy, which encourages the model to maximize the utilization of partial sequence information for predicting masked regions, rather than overfitting in highly redundant sequences. Moreover, the model’s training is more targeted, intensifying the focus on the highly diverse CDR-H3. This enables PARA to perform well in both FRs and CDRs, as well as multiple downstream tasks.

## Author contributions statement

L.L contributed conceptualization, project administration, review. X.G contributed conceptualization, methodology, validation, writing—review & editing. C.C contributed data curation, formal analysis, investigation, methodology, validation, writing.

